# Direct modulation of CRH nerve terminal function by noradrenaline and corticosterone

**DOI:** 10.1101/2024.06.11.598540

**Authors:** Emmet M. Power, Dharshini Ganeshan, Jamieson Paul, Hiroyuki Igarashi, Wataru Inoue, Karl J. Iremonger

**Affiliations:** Centre for Neuroendocrinology and Department of Physiology, School of Biomedical Sciences, University of Otago, Dunedin, New Zealand; Robarts Research Institute, Schulich School of Medicine and Dentistry, Western University, London, ON, N6A 5B7 Canada; Department of Pharmacology, Graduate School of Pharmaceutical Sciences, Tohoku University, Sendai, Miyagi, Japan

## Abstract

Nerve terminals are the final point of regulation before neurosecretion. As such, neuromodulators acting on nerve terminals can exert significant influence on neural signalling. Hypothalamic corticotropin-releasing hormone (CRH) neurons send axonal projections to the median eminence where CRH is secreted to stimulate the hypothalamic-pituitary-adrenal (HPA) axis. Noradrenaline and corticosterone are two of the most important neuromodulators of HPA axis function; noradrenaline excites CRH neurons and corticosterone inhibits CRH neurons by negative feedback. Here, we used GCaMP6f Ca^2+^ imaging and measurement of nerve terminal CRH secretion using sniffer cells to determine whether these neuromodulators act directly on CRH nerve terminals. Contrary to expectations, noradrenaline inhibited action potential-dependent Ca^2+^ elevations in CRH nerve terminals and suppressed evoked CRH secretion. This inhibitory effect was blocked by α2-adrenoreceptor antagonism. Corticosterone also suppressed evoked CRH peptide secretion from nerve terminals, independent of action potential-dependent Ca^2+^ levels. This inhibition was prevented by the glucocorticoid receptor antagonist, RU486, and indicates that CRH nerve terminals may be a site of fast glucocorticoid negative feedback. Together these findings establish median eminence nerve terminals as a key site for regulation of the HPA axis.

**Significance:** Corticotropin-releasing hormone (CRH) neurons control the stress axis. Noradrenaline and corticosterone are two signalling molecules that control CRH neuron cell body excitability. However, their effect on CRH nerve terminal function is unknown. To examine this, we performed live Ca^2+^ imaging and measured CRH secretion. We found that noradrenaline suppressed nerve terminal Ca^2+^ levels and inhibited nerve terminal CRH secretion. Corticosterone had no effect on nerve terminal Ca^2+^, but inhibited nerve terminal CRH secretion. This suggests that CRH nerve terminals may be a site of fast corticosteroid negative feedback. Together, these data demonstrate that CRH nerve terminals are a critical point of regulation in the control of the stress axis.

## Introduction

Corticotropin-releasing hormone (CRH) neurons are located in the paraventricular nucleus (PVN) of the hypothalamus and control the Hypothalamic-Pituitary-Adrenal (HPA) axis as well as behavioral responses to stress (Herman and Cullinan, 1997; Ulrich-Lai and Herman, 2009; Füzesi et al., 2016; Sterley et al., 2018; Kim et al., 2019a; Kim et al., 2019b; Daviu et al., 2020). CRH neuron responses to threats are highly plastic, allowing organisms to mount appropriate stress responses in different behavioral or physiological states (Kim et al., 2019b; Daviu et al., 2020; Fuzesi et al., 2023). When activated, CRH neurons release CRH from their nerve terminals in the median eminence, driving release of adrenocorticotropic hormone (ACTH) from the anterior pituitary gland. This in turn controls the release of corticosteroid stress hormones from the adrenal gland. Acute increases in corticosteroids lead to beneficial changes that promote survival after a stressful challenge. However, chronic elevations in corticosteroids can contribute to disease states (Lupien et al., 2009; Gourley et al., 2013). For this reason, CRH secretion must be kept under tight control.

CRH neurons are quickly activated when an animal encounters a real or potential threat (Füzesi et al., 2016; Kim et al., 2019a; Kim et al., 2019b; Daviu et al., 2020). Stress-dependent activation of CRH neurons is driven by afferent inputs, with noradrenergic inputs being especially important (Plotsky et al., 1989; Pacak et al., 1992; Ritter et al., 2003; Flak et al., 2014). We have recently shown that noradrenaline is a potent activator of CRH neurons, switching these neurons into a bursting mode of activity. This excitatory effect was mediated by α1-adrenoreceptors, with α2-adrenoreceptors limiting this excitation (Gouws et al., 2022). Noradrenergic fibers innervate PVN CRH neurons (Palkovits et al., 1980; Sawchenko and Swanson, 1982; Cunningham and Sawchenko, 1988; Dunn et al., 2004). Interestingly, noradrenergic fibers also innervate the median eminence where the CRH nerve terminals reside (Bjorklund et al., 1970; Jonsson et al., 1972; Palkovits et al., 1977; Palkovits et al., 1980). This raises the possibility that noradrenaline might also directly excite CRH nerve terminals in the median eminence.

To prevent excessive corticosteroid secretion, feedback mechanisms exist to suppress CRH neuron activity once corticosteroid levels become elevated (Bittar et al., 2019). This negative feedback is primarily mediated by glucocorticoid receptors (GRs) (Sapolsky et al., 2000; Gjerstad et al., 2018). GRs are present at high levels in CRH neurons and when activated, can suppress excitability by either fast non-genomic (Di et al., 2003; Nahar et al., 2015) or slow genomic pathways (Tasker and Herman, 2011; Kim et al., 2019b). Recent in vivo recordings of CRH neuron activity have demonstrated that glucocorticoid negative feedback is evident 30-40 minutes following elevation of corticosterone levels (Kim et al., 2019b). Importantly, these recordings were performed in the PVN where the CRH neuron cell bodies reside. This leaves open the possibility that fast forms of glucocorticoid negative feedback may occur in other parts of the CRH neuron, such as at the nerve terminals in the median eminence (Sarrieau et al., 1988).

Here, we set out to determine whether CRH nerve terminals are an independent point of HPA axis regulation by noradrenaline and corticosterone. To study median eminence nerve terminal function, we used a combination of in vitro GCaMP6f Ca^2+^ imaging and CRH receptor sniffer cells. corticosterone had no effect on action potential evoked Ca^2+^ levels in CRH nerve terminals. Nevertheless, corticosterone quickly suppressed CRH nerve terminal peptide secretion. This inhibition was mediated by GRs. Surprisingly, noradrenaline decreased action potential evoked nerve terminal Ca^2+^ levels and also inhibited CRH peptide release from median eminence nerve terminals. This inhibition was mediated by α2-adrenoreceptors. Together these data reveal that CRH nerve terminals in the median eminence are an independent point of regulation of CRH peptide secretion. These data also show that neuromodulators can exert different effects on excitability depending on the region of the neuron targeted.

## Methods

### Animals

All experiments were carried out in adult male (2-6 months old) *Crh-Ires-cre x Ai148-GCaMP6f* or C57BL6J mice. Animals had a 12h light/dark cycle (7am-7pm lights on) with food (2918 Teklad Irradiated Global 18% Protein Rodent Diet, Inotiv) and water available *ad libitum*. All protocols and procedures were approved by the University of Otago Animal Ethics Committee and carried out in accordance with the New Zealand Animal Welfare Act.

### Immunohistochemistry

Paraformaldehyde fixed coronal brain sections (30µm thick) of the median eminence were labelled for GCaMP6f with a GFP antibody (chicken anti-GFP; 1:2000; Aves Labs) and for CRH peptide (polyclonal rabbit anti-CRH, 1:1000, Abcam). Secondary antibodies used for visualisation were goat anti-chicken IgG Alexa Fluor-488 (1:2000, Thermo Fisher Scientific) and goat anti-rabbit IgG Alexa Fluor-567 (1:1000, Thermo Fisher Scientific). Sections were imaged under an Olympus BX51 fluorescence microscope.

### Brain slice preparation

Mice were killed by cervical dislocation between 9am and 10:30am, their brain quickly removed and placed in ice-cold oxygenated (carbogen 95% O_2_, 5% CO_2_) slicing solution containing (in mM); 87 NaCl, 2.5 KCl, 25 NaHCO_3_, 1.25 NaH_2_PO_4_, 0.5 CaCl_2_, 6 MgCl_2_, 25 D-Glucose, 75 sucrose, pH 7.2-7.4. 200 µm thick coronal slices of the median eminence were then cut using a vibratome (VT1200S, Lecia Microsystems). These slices were then incubated in a holding chamber in oxygenated artificial cerebrospinal fluid (ACSF) containing in (mM); 126 NaCl, 2.5 KCl, 26 NaHCO_3_, 1.25 NaH_2_PO_4_, 2.5 CaCl_2_, 1.5 MgCl_2_, 10 D-Glucose at 30°C for at least 1 hour before recording.

### Electrical stimulation and GCaMP6f Ca^2+^ imaging

Brain slices containing the median eminence were secured in a tissue bath and constantly perfused with carbogen-bubbled warm (30°C) ACSF at a rate of 1-2 ml/min. Extracellular electrical stimulation was delivered using a monopolar borosilicate glass electrode filled with ACSF and positioned in the internal zone of the median eminence. Biphasic train stimulations, between 0.1-0.5 µA, were delivered using a S88X GRASS stimulator (Astro-med inc) at 10 Hz for two seconds. Imaging was performed with an Olympus FV1000 confocal microscope fitted with an Olympus 40X, 0.8 NA objective lens. GCaMP6f was excited with a 488 nm Argon laser (Melles Griot). Emitted light was detected by a PMT after passing through a bandpass filter (505-605 nm). The confocal aperture was wide open during Ca^2+^ imaging experiments to collect maximum emitted fluorescence. Frame scans were performed on zoomed regions at approximately 2Hz. Images were captured using Fluoview 1000 software and analyzed using ImageJ. Changes in fluorescence were calculated as ΔF/F, where F is the averaged (pre-stimulation) baseline fluorescence for each region of interest (ROI). All analysis compared peak ΔF/F values for each ROI, identified as the maximum value reached during stimulation. ROIs were identified manually in response to the first stimulation and were consistent across all subsequent timepoints. ROIs consisted of clusters of boutons and were always located in the external zone of the median eminence. ROIs were deemed unstable and excluded from analysis if GCaMP6f fluorescence responses at the 25 min timepoint (the last baseline timepoint) deviated by more than 15% compared to an average of all previous baseline responses for that ROI. Data was normalized to an average of all baseline values (i.e. timepoints 0-25 min).

### Sniffer cell culture and sniffer cell application to brain slices

HEK-293 cells stably expressing pCMV-CRHR1-mRFP/pCAGGS-mTurq-EPAC-citrine were used to detect CRH secretion and are hence referred to as sniffer cells. Sniffer cells expressed both the CRHR1 receptor and an EPAC-based FRET sensor. The FRET sensor consisted of variants of Cyan-Fluorescent-protein (CFP) and Yellow-Fluorescent-protein (YFP, Fig 4A). When CRH binds to the CRHR1 on sniffer cells, there is an increase in cAMP which leads to a confirmational change in the EPAC-FRET sensor. This causes a reduction in FRET, i.e. CFP emission increases and YFP emission decreases (Fig 4Aiii).

Cell culture supplies were purchased from Life Technologies NZ Ltd., unless otherwise stated. Sniffer cells were cultured in high glucose DMEM growth medium containing 1% sodium pyruvate, 1% Pen/strep, 1.2% geneticin and 10% fetal bovine serum (FBS, Hyclone, Sigma Aldrich). Cells were incubated at 37°C and 5% CO_2_, until an 80% confluent monolayer (average cell doubling time 12.5 hrs) was reached with a typical morphology of epithelial-like cells.

Adherent cells were initially washed with DPBS and were detached by incubating with trypsin EDTA (0.25%) at 37°C. Subsequently, cells were neutralised with 10% FBS growth medium, subsampled for cell count and centrifuged at 1000 rpm for 5 min. After centrifugation, the supernatant was discarded, and the cells were reconstituted in warm ACSF with a final concentration of 10,000 cells/µl to be applied to median eminence brain sections. For experiments imaging sniffer cells adhered to glass coverslips, the cells were reconstituted into growth medium and plated at 5000 cells/cm2 on 13 mm borosilicate glass coverslips (Trajan, Cat no. 471111300), pretreated with poly-L-lysine (Sigma, P4707). Cells were given 48 hours to grow before imaging.

For experiments involving sniffer cell responses to median eminence stimulation, brain slices were prepared as outlined above. 5-15 minutes following slice preparation, sniffer cells suspended in ACSF were pipetted onto the median eminence region of brain slices maintained within the normal incubation chamber. Each slice received ̴100 µl at 10,000 cells/µl concentration. Once applied slices were given at least 1 hour to allow sniffer cell adhesion before being transferred to a recording chamber where they were continually perfused with ACSF.

### Sniffer cell imaging

Sniffer cell imaging was performed on an Olympus upright epifluorescence microscope connected to a pE300 CoolLED light source with a 40X (0.8 NA) water dipping objective. CFP was excited with blue (434nm peak) light using a narrow violet dichroic filter (U-MNV2, Nikon). Images were captured at 2 Hz with a Prime BSI camera (Teledyne Photometrics, USA) fitted to the microscope via an Optosplit II (Cairn) emission image splitter containing 455nm long-pass, 475nm CFP and 527nm YFP emission filters. This allowed for the simultaneous capture of both CFP and YFP channels.

Sniffer cell images were analysed using ImageJ and the Cairn image splitter plugin. Regions of interest (ROI) were defined as individual sniffer cells, or pairs of cells when indistinguishable. For median eminence brain slice experiments, only sniffer cells adhered to the median eminence were identified as ROIs. Sniffer cells adhered to third ventricle or elsewhere were not analyzed. Ratiometric fluorescence responses were calculated by first calculating a ratio response by dividing the CFP signal by the YFP signal. This ratio response was then used to calculate ΔF/F, where F is an average of the baseline period ratio and ΔF is the peak change in ratio after CRH/stimulation. Fluorescence responses were bleach corrected using no stimulation control recordings. ROIs that did not exceed a peak of 2% ΔF/F response were not included.

### Stimulation and drug application protocols

For sniffer cells applied to median eminence slices, the first electrical stimulation evoked the largest response, which was consistently reduced in amplitude on a subsequent stimulation. However, we found that the 2^nd^ and 3^rd^ stimulations evoked responses which were of consistent size. Therefore, the first stimulation was collected but not analyzed (except in the case of Astressin experiments, mentioned below). All experiments reported here compared the second and third timepoints of stimulation responses, which had a 20-minute time interval between them.

For slice experiments all drug applications had approximately 15-minute wash on before recordings began, with the exception of Astressin which required pre-incubation of the slice for approximately one hour before recording. As Astressin required pre-incubation, repeated timepoint experiments were not possible. We therefore compared the first sniffer cell response (to CRH peptide or terminal stimulation) in the presence of Astressin to the first response of sniffer cells in control conditions (ie no Astressin).

To evoke CRH peptide release from median eminence nerve terminals, a monopolar glass electrode was positioned in the internal zone of the median eminence, similar to GCaMP6f recordings. Stimulation consisted of biphasic pulses at 0.3 µA (3 Hz for 3 seconds). Sniffer cell imaging was 5 minutes long and stimulation occurred at 30 seconds after the start of recording. Repeat recordings were performed at 20-minute intervals.

For coverslip experiments sniffer cells in the recording chamber were constantly perfused with ACSF at 2 ml/min. Bath application of CRH peptide was done via the ACSF perfusion system. For control recordings, CRH dissolved in ACSF (1 nM, Sigma-Aldrich) was introduced to the perfusion line for 50 seconds (except for Figure 4c when it was applied for 120 seconds). When applied in combination with other drugs (e.g. noradrenaline), these drugs were also included in the ACSF+CRH peptide solution and applied for 50 seconds. However, all other drugs were in the ACSF flow prior to (15 min) and after CRH application at the same concentrations. For analysis of both coverslip and slice sniffer cell imaging, peak ratiometric ΔF/F values were used, measured as the maximum ΔF/F value reached throughout the entire recording for each individual ROI.

### Statistical analysis

Statistical analysis was performed using GraphPad Prism 10. All reported values are the mean ± SEM. Comparisons between groups were carried out using student T-tests, One or Two-way ANOVA where appropriate, with Tukey’s or Sidak’s post hoc multiple comparison tests. ROI number is reported as “n” and slice number (or coverslip number) is reported as “N”. Three or more mice were used per experiment. P<0.05 was considered statistically significant.

## Results

### Characterisation of stimulation-evoked Ca^2+^ responses in CRH nerve terminals

To study the regulation of CRH neuron nerve terminals in the median eminence, we used *CRH-ires-cre x Ai148-GCaMP6f* mice. Immunostaining for CRH peptide and GCaMP6f (GFP) demonstrated colocalization predominantly in the external zone of the median eminence, the region of CRH peptide storage and release (Fig 1A).

**Figure 1.**
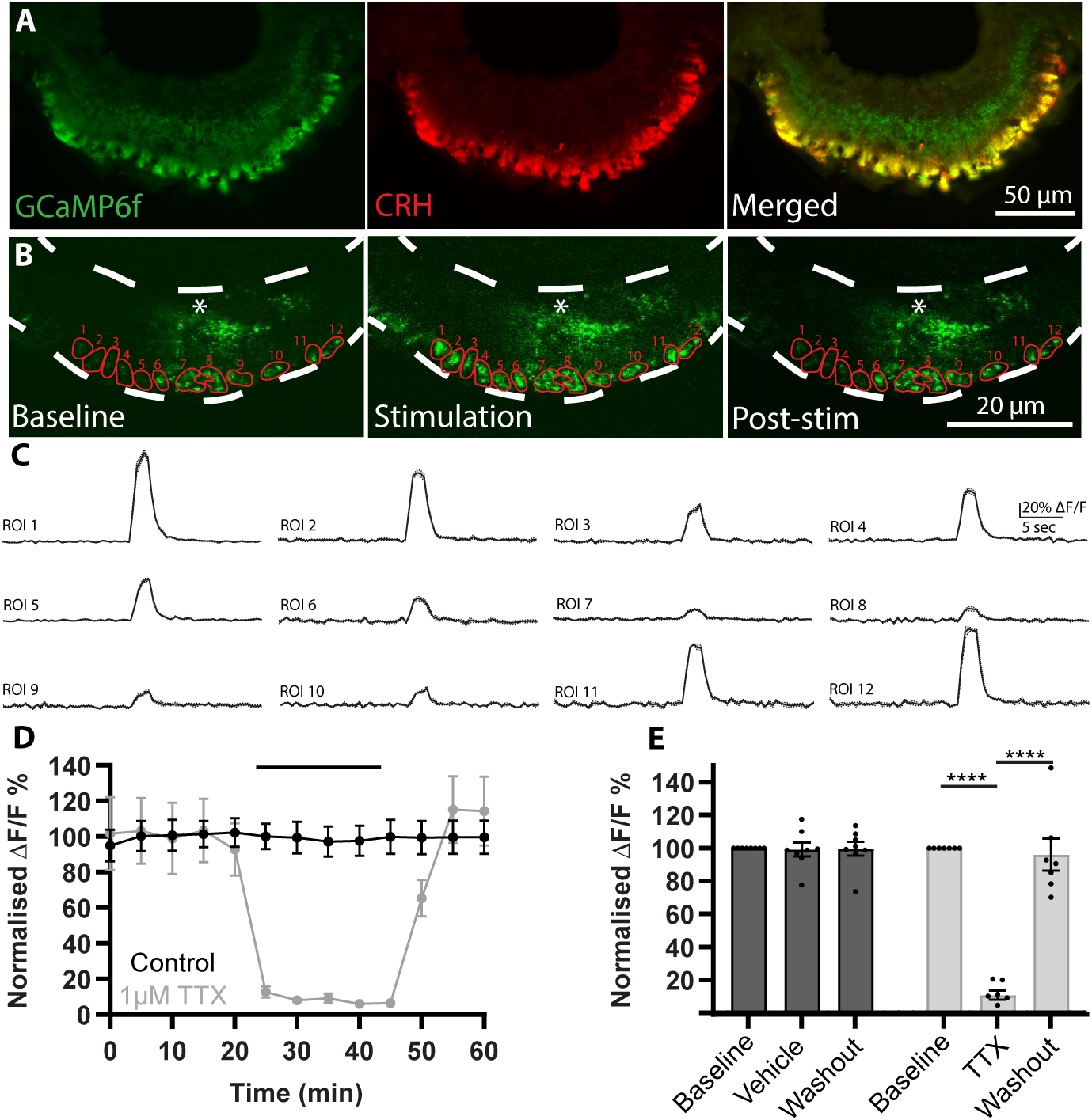
GCaMP6f Ca^2+^ imaging of CRH nerve terminals. **A.** Images of the median eminence stained for GCaMP6f (left, green), CRH peptide (middle, red) and the merged image (right). **B.** Single frame captures showing GCaMP6f fluorescence in the median eminence pre-stimulation (baseline), during stimulation and post-stimulation (post-stim). Red numbered shapes denote regions of interest (ROIs). White asterisks indicates position of stimulating electrode. **C.** Example fluorescence responses from the ROIs identified in B. **D.** Peak amplitude of GCaMP6f responses plotted over time for control (black, N=3, n=8) and 1 µM TTX (grey, N=3, n=7) treated slices. Black horizontal bar indicates the time of TTX wash in. **E.** Data from (D) has been divided into baseline (period prior to TTX wash on, 20 min), treatment (vehicle or TTX wash on period, 20 min) and wash off periods (post wash on period, 20 min). TTX induced a significant inhibition of the peak GCaMP6f response (*P<0.05, ***P<0.001, Post hoc Sidak’s multiple comparisons test).

To evoke action potentials in CRH neuron axons, a stimulating electrode was placed in the internal zone of the median eminence. Bursts of stimulation (10 Hz, 2 sec) evoked transient increases in GCaMP6f fluorescence in CRH terminal boutons in the external zone (Fig 1B). While stimulation-evoked responses varied in amplitude across the median eminence, the kinetics of responses were similar (Fig 1C). When stimulated once every 5 minutes, evoked responses were stable over a 60-minute recording period (Fig 1D/E). To confirm these evoked Ca^2+^ responses were action potential dependent, we blocked voltage gated Na^+^ channels using tetrodotoxin (TTX 1 µM, Alomone labs). This resulted in a robust but reversible inhibition of the evoked response (Fig 1D/E, Two-way ANOVA, Treatment: F_(2,26)_=42.95, P<0.0001, Time: F_(2,26)_=55.07, P<0.0001, Interaction: F_(1,13)_=11.38, P=0.005).

### Corticosterone does not change CRH nerve terminal Ca^2+^ responses

Corticosterone acts to inhibit glutamate synaptic input (Di et al., 2003; Nahar et al., 2015) and the excitability of CRH neurons when measured at the cell soma in the PVN (Senst et al., 2016; Kim et al., 2019b). To test if corticosterone can also inhibit CRH nerve terminals at the level of the median eminence, we measured stimulation-evoked Ca^2+^ responses before, during and after bath application of 1 µM corticosterone(Sigma-Aldrich). Interestingly, corticosterone application did not affect stimulation-evoked fluorescence responses in CRH terminals (Fig 2A/B). Comparison of corticosterone-treated and control groups showed no significant difference between the two (Fig 2C, Two-way ANOVA, Treatment: F_(1,294)_=0.105, P=0.747, Time: F_(2,294)_=0.097, P=0.908, Interaction: F_(2,294)_=0.026, P=0.974).

**Figure 2.**
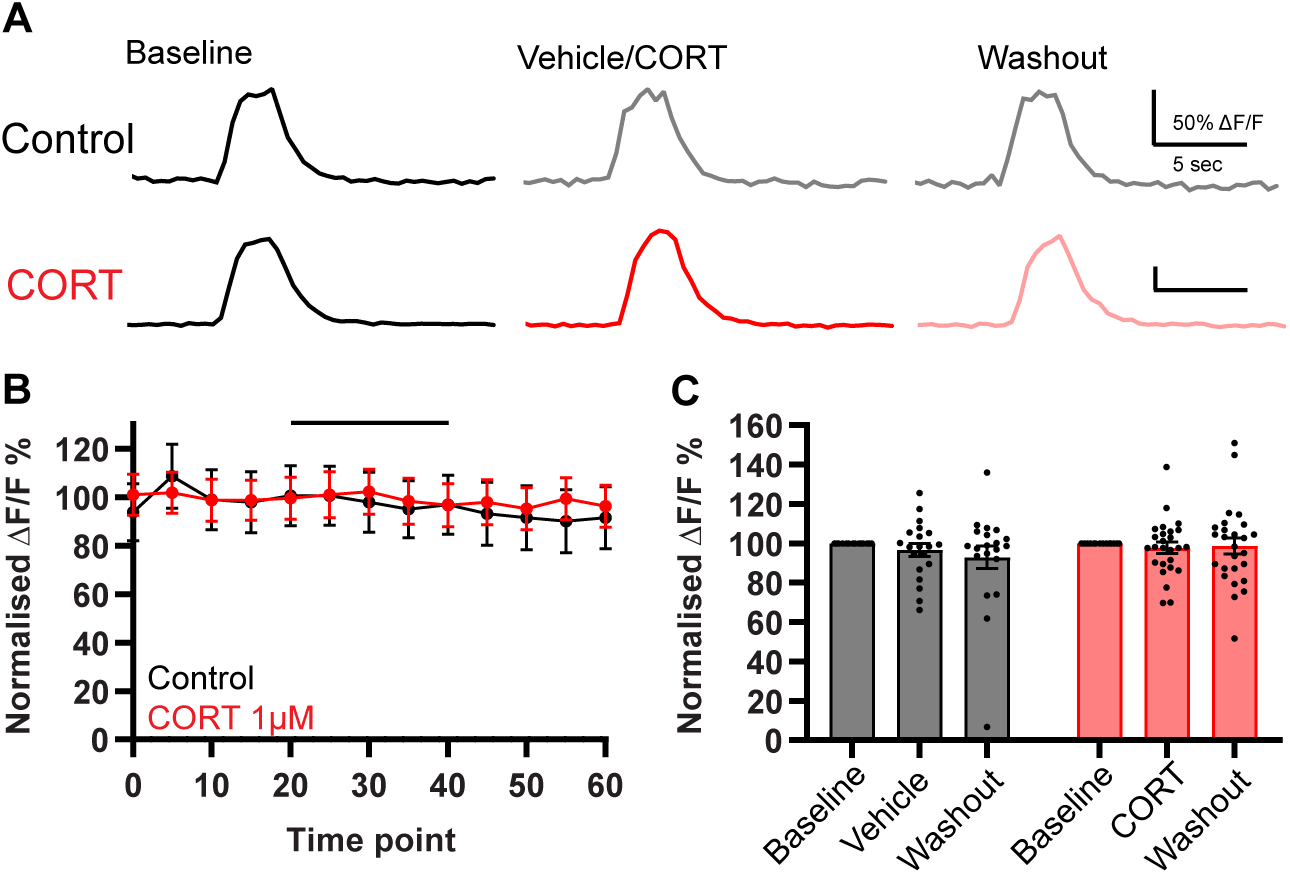
Corticosterone does not regulate CRH nerve terminal Ca^2+^ responses. **A.** Representative GCaMP6f fluorescence responses to electrical stimulation from Control and corticosterone (CORT) treated groups. **B.** Peak GCaMP6f responses from control recordings (black, N=4, n=20) and those where 1 µM CORT was washed in (red, N=4, n=26). Black horizontal bar indicates the time of CORT or vehicle wash in. **C.** Bar graph of data showing averaged peak responses for baseline, treatment and wash off periods.

### Noradrenaline inhibits CRH nerve terminal Ca^2+^ responses

We next tested the effect of a neuromodulator previously shown to have excitatory actions on CRH neuron cell bodies in the PVN (Gouws et al., 2022). To examine the effect of noradrenaline on Ca^2+^ responses in the median eminence we measured stimulation-evoked Ca^2+^ responses before, during, and after bath application of noradrenaline. Noradrenaline (Sigma-Aldrich), at either 10, 20, or 100 µM, significantly inhibited stimulation-evoked Ca^2+^ responses in CRH nerve terminals (Fig 3A/B). 100 µM noradrenaline decreased peak Ca^2+^ response by an average of 46.79 ± 4.8% compared to baseline. 20µM noradrenaline had a similar effect decreasing peak responses by 48.9 ± 6.13% compared to baseline, while 10 µM had a slightly less potent effect decreasing responses by 28.97 ± 4.68%.

**Figure 3.**
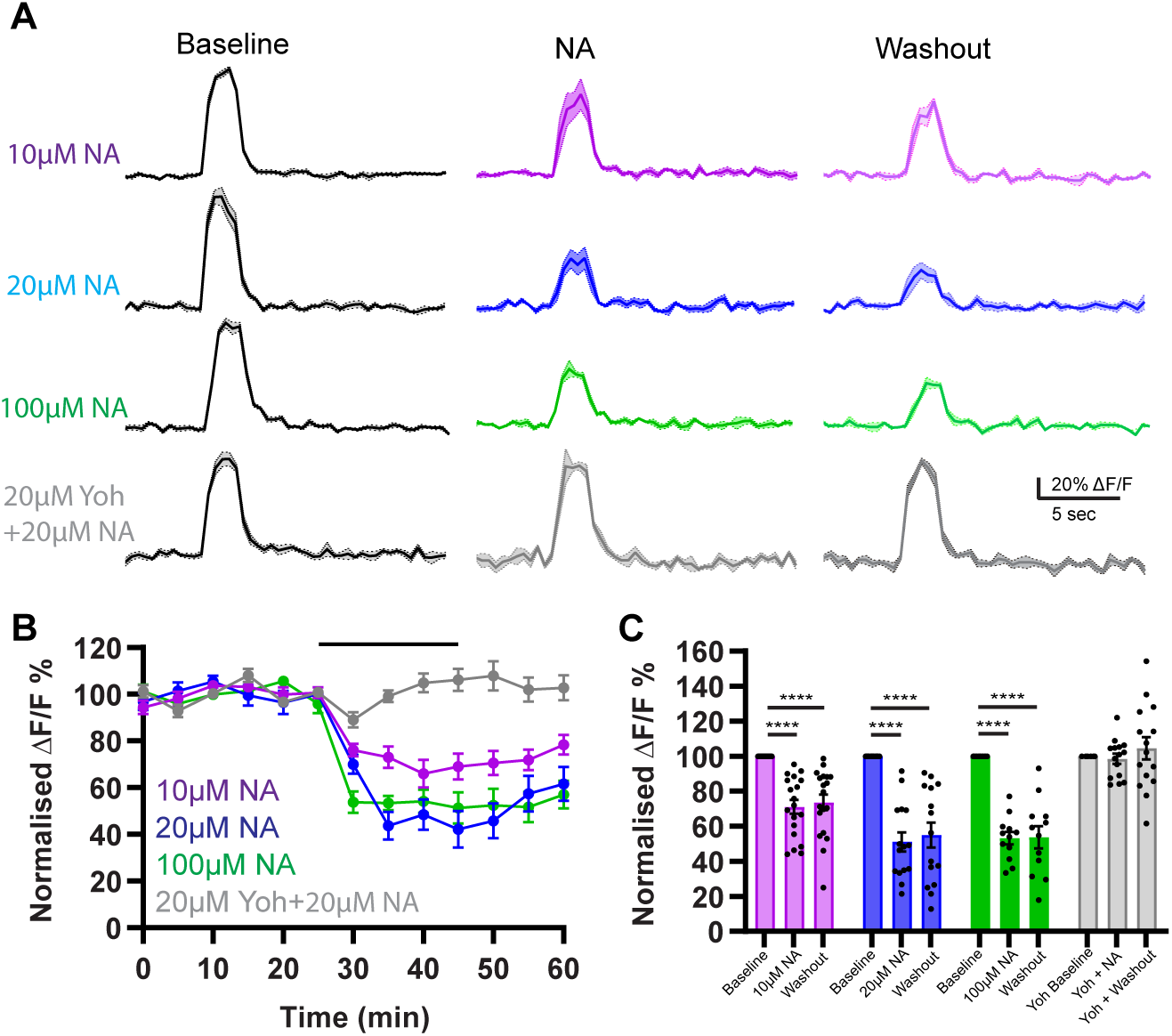
Noradrenaline (NA) inhibits CRH nerve terminal Ca^2+^ responses. **A.** Mean ± SEM GCaMP6f fluorescence responses to electrical stimulation before and after bath application of noradrenaline at different concentrations. **B.** Peak amplitude of GCaMP6f responses plotted over time for the various treatment groups. Black line denotes time of treatment. Colours correspond to treatment received and match traces in (A). **C.** Summary data comparing the baseline (first 25 min), treatment (following 20 min) and wash off periods (final 15 min) for each group. A Two-way ANOVA revealed a significant difference between the groups (See Results section) with Sidak’s multiple comparison test results denoted by asterisks on the graph. ****P<0.0001. 10 µM NA: N=4, n=19, 20 µM NA: N=3, n=15, 100 µM NA: N=3, n=12, 20 µM Yoh+20 µM NA: N=3, n=18.

Previous work suggests that CRH neurons express the α2-adrenoreceptor (Chen et al., 2019; Gouws et al., 2022). To determine if the α2-adrenoreceptor was responsible for the inhibition of Ca^2+^ responses in CRH nerve terminals, noradrenaline (20 µM) was applied in the presence of yohimbine (20 µM, Sigma-Aldrich), an α2-adrenoreceptor antagonist. In the presence of yohimbine, noradrenaline no longer inhibited stimulation-evoked Ca^2+^ responses (Fig 3 A/B). Statistical comparison of the groups (broken into baseline, treatment and wash off periods) showed a significant difference between the treatments (Fig 3C, Two-way ANOVA, Treatment: F_(4,249)_=34.34, P<0.0001, Time: F_(2,2,249)_=38.92, P<0.0001, Interaction: F_(8,249)_=8.7, P.0001). Taken together, these data reveal that noradrenaline inhibits stimulation-evoked Ca^2+^ responses in CRH nerve terminals in the median eminence, despite its excitatory effects on CRH soma within the PVN (Gouws et al., 2022).

### CRHR1 sniffer cells can reliably detect CRH peptide

We next set out to determine if corticosterone or noradrenaline could regulate CRH peptide secretion from nerve terminals in the median eminence. To measure CRH secretion, we used CRHR1 sniffer cells (Fig 4A, see methods). To validate the sniffer cell responses, coverslips with sniffer cells were imaged before, during and after bath application of CRH peptide (1 nM, Sigma-Aldrich, 50 sec, Fig 4Aiii). CRH induced a marked change in the ΔF/F FRET ratio that returned to baseline upon washout. To test the effects of drugs on sniffer cell responses to CRH peptide, we used a two-exposure protocol, where the first exposure was used as control and the second as a treatment timepoint. There was no significant difference in the response to bath applied CRH using this two-exposure protocol (Fig 4B, Paired T-test, P=0.673). We next tested if the sniffer cell responses were blocked with the CRHR1 antagonist Astressin. Sniffer cells needed to be incubated in Astressin (500 nM, Tocris) for 10+ mins in order to fully block CRHR1 receptors. After incubation, CRH peptide was washed on and responses were compared to control cells exposed to CRH peptide alone. Astressin significantly decreased the fluorescence response of the sniffer cells to CRH application (Fig 4C, Unpaired T-test, P<0.0001).

**Figure 4.**
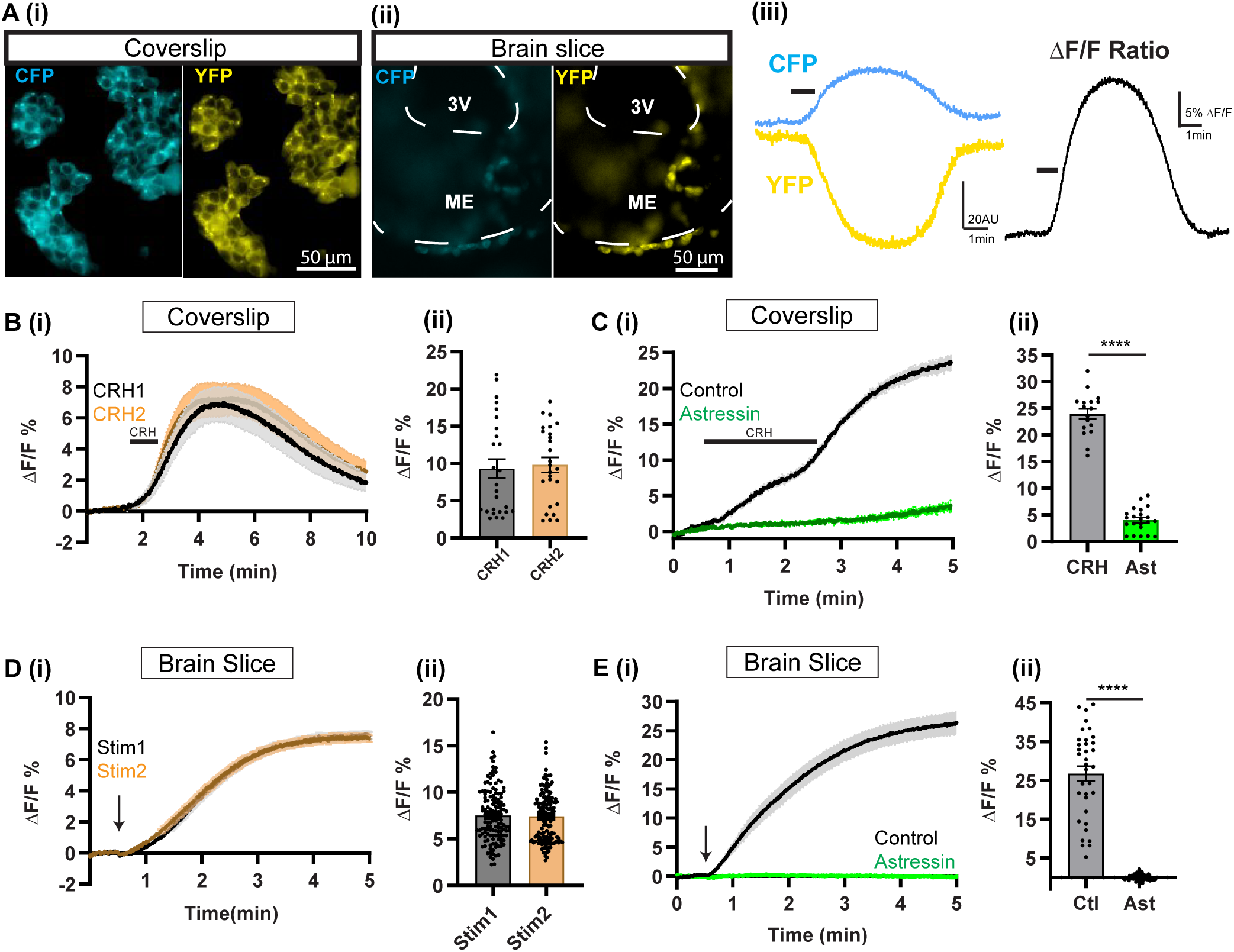
CRHR1 sniffer cells reliably detect CRH peptide. **A (i)** Images of sniffer cells cultured on a glass coverslip (both CFP and YFP channels shown). **(ii)** Image of sniffer cells adhered to a coronal section of the median eminence (ME). The white dotted lines denote the internal and external limits of the median eminence. 3V = third ventricle. **(iii)** Example response from sniffer cells cultured on a glass coverslip in response to 50 sec bath application of 1 nM CRH. Wash on time denoted by black bar. Ratio-metric (CFP/YFP) response was used to calculate ΔF/F, black trace on right. All subsequent data shown is the ratiometric ΔF/F response. **B (i)** Averaged response of the sniffer cells, on coverslips, to two consecutive exposures to CRH peptide (1 nM, 50 sec each). Black trace is the response to first exposure, brown trace is the response to second exposure, N=5 covers-lips, n=26. **(ii)** Summary data showing no difference in peak responses between the two CRH applications. **C (i)** Sniffer cell responses to CRH peptide in the presence (green, N=3, n=22) and absence (black, N=3 coverslips, n=18) of CRHR1 antagonist Astressin (Ast, 0.5 µM). **(ii)** Peak ΔF/F responses were significantly lower in the presence of Astressin. **D (i)** Averaged sniffer cell responses to two consecutive electrical stimulations of the median eminence. Arrow indicates time of stimulation (N=7, n=137). **(ii)** Summary data showing no significant difference in peak ΔF/F responses between stim 1 and stim 2. **E (i)** Sniffer cells responses to median eminence stimulation in control (N=5, n=35) and Astressin (0.5 µM) treated slices (N=4, n=82). **(ii)** Peak responses of sniffer cells were blocked in the presence of Astressin. Asterisks indicate significant results of Unpaired T-test (see Results) ****P<0.0001.

For *in vitro* slice experiments, suspended CRHR1 sniffer cells were applied onto the median eminence region of brain slices and imaged (Fig 4Aii). A two stimulation protocol was then used to validate the sniffer cell responses on slices. To induce endogenous CRH release, we stimulated the median eminence for 3 sec at 3 Hz, with 20 minutes given between stimulations. There was no significant difference in the sniffer cell response between the first and second stimulations (Fig 4D, Paired T-test, P=0.557). To confirm that the sniffer cell responses were due to release of CRH peptide, we incubated slices in Astressin (500 nM) prior to stimulation. This significantly inhibited the sniffer cell response to median eminence stimulation compared to control slices (Fig 4E, Unpaired T-test, P<0.0001). Together, these data show that CRHR1 sniffer cells can be used to detect secreted CRH peptide from median eminence brain slices.

### Corticosterone and noradrenaline both decrease sniffer cell responses to median eminence stimulation *in vitro*

While we found that corticosterone had no effect on stimulation-evoked Ca^2+^ responses in CRH nerve terminals, the possibility remained that corticosterone could inhibit CRH release via a Ca^2+^-independent pathway. To ensure corticosterone had no direct effect on the sniffer cells, we first ran a control experiment on cells plated on coverslips. As in the previous coverslip experiment the cells were exposed to two brief (50 sec) applications of CRH peptide. In this case however, the second application was in the presence of corticosterone (1 µM). Corticosterone had no significant effect on sniffer cell responses to CRH peptide (Fig 5A, Paired T-test, P=0.544).

**Figure 5.**
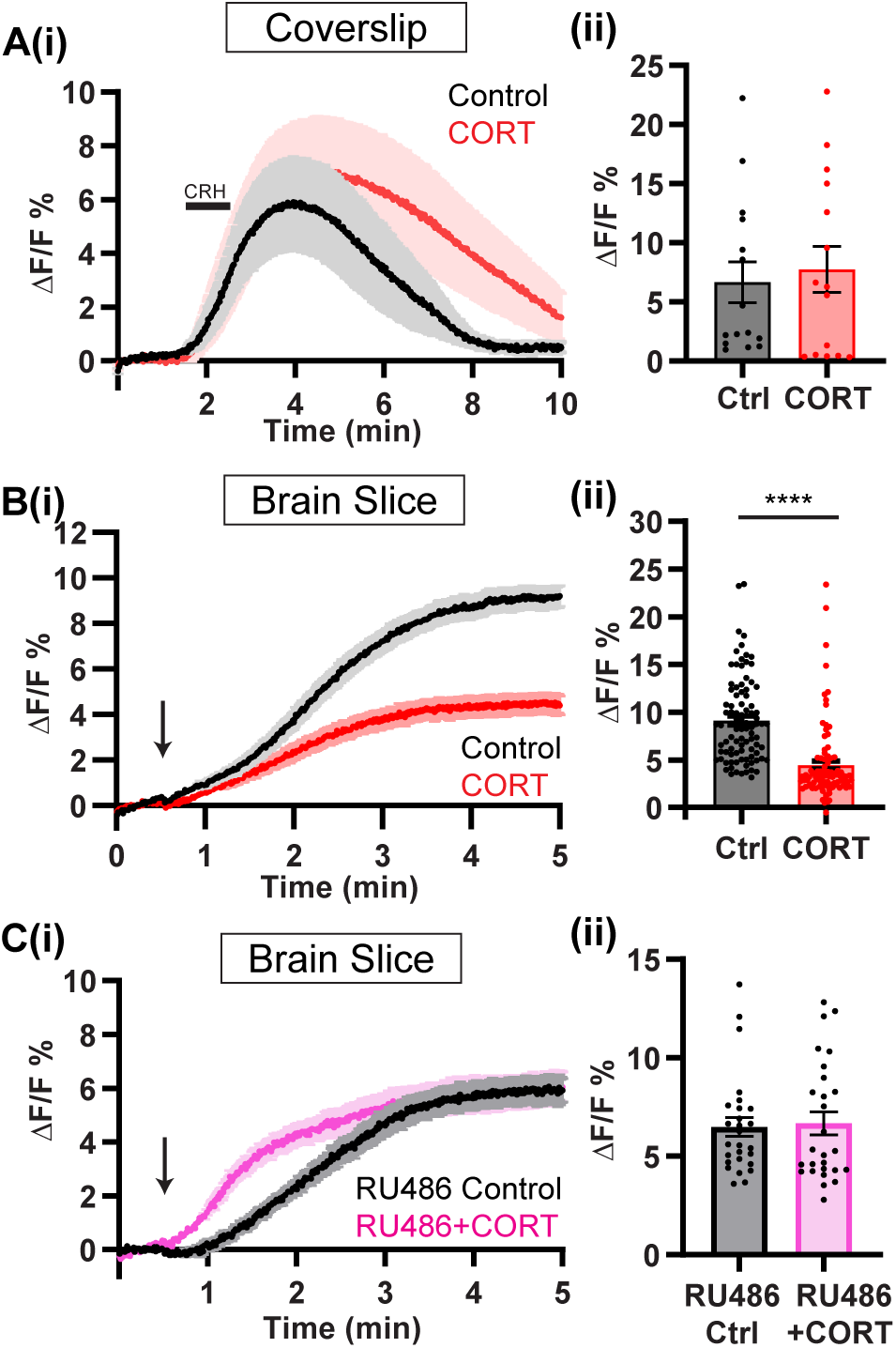
Corticosterone inhibits sniffer cell responses to median eminence stimulation. **A (i)** Response of sniffer cells (on coverslips) to bath applied CRH peptide before (black) and after (red) the bath application of corticosterone (CORT, 1 µM, N=4 coverslips, n=15). **(ii)** Peak ΔF/F responses to CRH peptide were not significantly different before (control) and after CORT treatment. **B (i)** Response of sniffer cells (adhered to brain slices) to median eminence stimulation before and after CORT. Arrow indicates time of stimulation. N=6, n=91 **(ii)** Peak ΔF/F responses to median eminence stimulation are significantly reduced after application of CORT. ****P<0.0001. **C (i)** RU486 was applied to block GRs and then sniffer cells responses to median eminence stimulation were determined before and after application of CORT (N=3, n=27). **(ii)** In the presence of RU486, peak ΔF/F responses to median eminence stimulation are not significantly changed by CORT.

We then tested the effect of corticosterone on CRH peptide release from the median eminence using the two-stimulation protocol in brain slices. Interestingly, we found that corticosterone induced a significant reduction in sniffer cell responses when compared to control stimulation (Fig 5B, Paired T-test, P<0.0001). We next tested if this inhibition could be prevented with the glucocorticoid receptor antagonist RU486 (1 µM, Tocris). In the presence of RU486, corticosterone no longer inhibited sniffer cell responses to median eminence stimulation (Fig 5C, Paired T-test, P=0.754). These data together suggest corticosterone is acting via the glucocorticoid receptor to induce a Ca^2+^-independent suppression of secretion in CRH nerve terminals.

Next, we investigated if noradrenaline was having a similar effect on CRH peptide release. HEK293 cells endogenously express α1-adrenoreceptors (Atwood et al., 2011; Pillay et al., 2020), and control experiments showed they were significantly excited by the application of noradrenaline, but this excitation was blocked by α1-adrenoreceptor antagonist prazosin (T-test peak ΔF/F response, P<0.0001, data not shown). For this reason, all sniffer imaging with noradrenaline was performed in the continued presence of prazosin (10 µM, Sigma-Aldrich). We first tested if noradrenaline had any effect on sniffer cell responses following bath application of CRH peptide. In the presence of prazosin, noradrenaline had no significant effect on sniffer cell responses to CRH peptide (Fig 6A, Paired T-test, P=0.176). However, application of noradrenaline to *in vitro* slices did decrease the stimulation-evoked release of CRH peptide as measured by sniffer cell responses (Fig 6B, Paired T-test, P<0.0001). This inhibitory effect of noradrenaline was blocked with the α2-adrenoreceptor antagonist yohimbine (20 µM) (Fig 6C, Paired T-test, P=0.11). These data confirm that noradrenaline suppresses CRH peptide release from median eminence nerve terminals via α2-adrenoreceptors.

**Figure 6.**
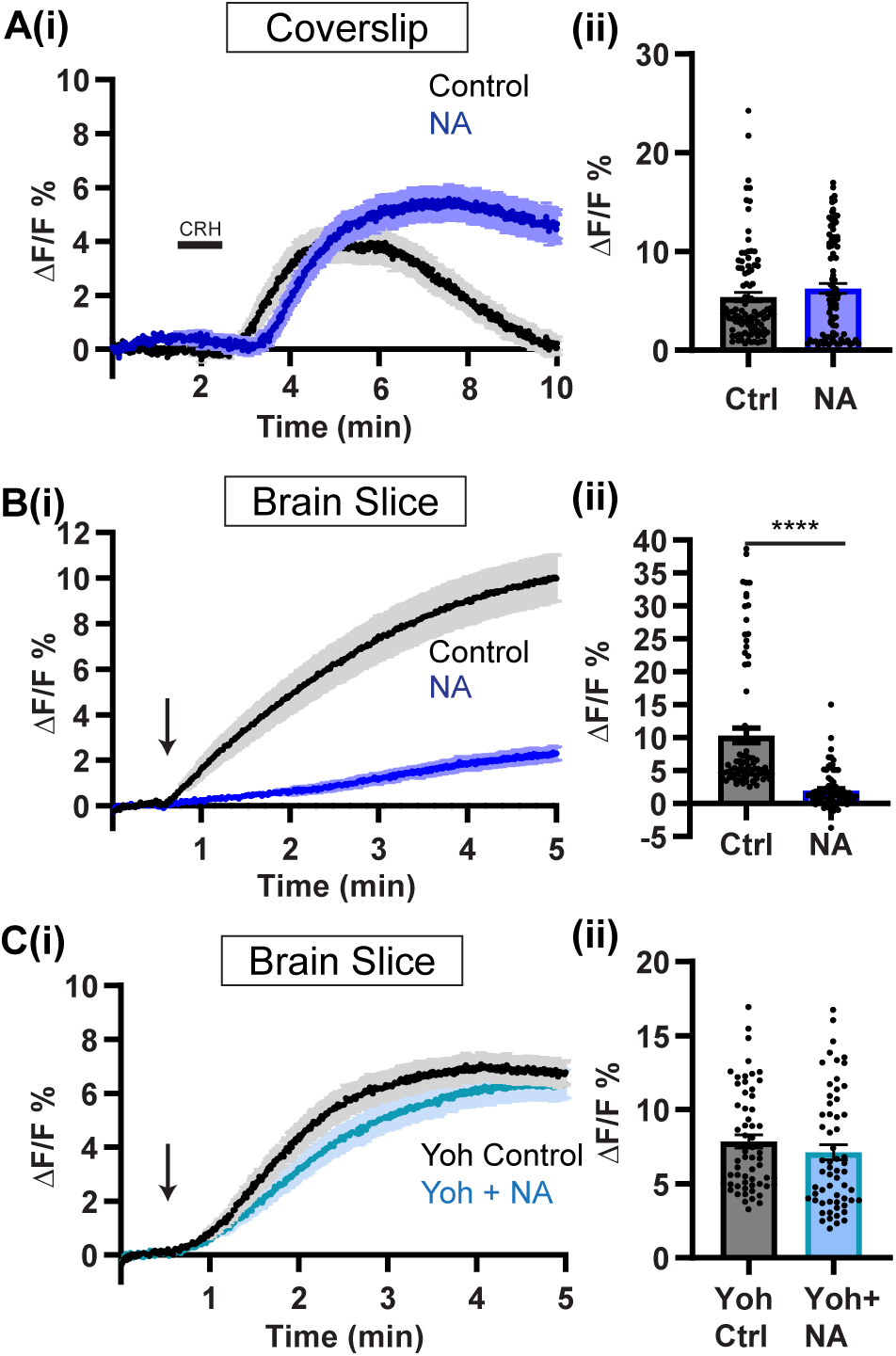
Noradrenaline inhibits sniffer cell responses to median eminence stimulation. **A (i)** Response of sniffer cells (on coverslips) to bath applied CRH peptide before (black) and after (blue) the bath application of noradrenaline (20 µM; N=4 coverslips, n=97). All experiments in Figure 6 were performed in the presence of prazosin (10 µM). **(ii)** Peak ΔF/F responses to CRH peptide were not significantly different before (control) and after noradrenaline treatment. **B (i)** Response of sniffer cells (adhered to brain slices) to median eminence stimulation before and after noradrenaline (N=6, n=86). Arrow indicates time of stimulation. **(ii)** Peak ΔF/F responses to median eminence stimulation are significantly reduced after application of noradrenaline. ****P<0.0001. **C (i)** Yohimbine (Yoh, 20 µM) was applied to block α2-adrenoreceptors and then sniffer cells responses to median eminence stimulation were determined before and after application of noradrenaline (N=4, n=60). **(ii)** In the presence of yohimbine, peak ΔF/F responses to median eminence stimulation are not significantly changed by CORT.

## Discussion

While neuromodulators play and important role in regulating the activity of CRH neurons and hence the HPA axis, studies to date have focused on regulation of CRH neuron excitability at the cell body (Hoyda et al., 2009; Inoue et al., 2013; Nahar et al., 2015; Jamieson et al., 2017; Pati et al., 2020). Here, we show that two critical neuromodulators that regulate stress responses also exert powerful actions directly on CRH nerve terminals in the median eminence. Specifically, we show that noradrenaline inhibits action potential dependent Ca^2+^ elevations in CRH nerve terminals and suppresses CRH peptide secretion. This effect is the opposite of noradrenaline’s well-established excitatory action at CRH neuron cell bodies (Chen et al., 2019; Gouws et al., 2022). Corticosterone also suppressed evoked CRH peptide secretion, but this occurred without any change in action potential dependent Ca^2+^. Together, these data reveal that the CRH nerve terminal is a site for neuromodulation of CRH secretion and regulation of the stress axis.

PVN CRH neurons receive noradrenergic projections from the A1 cell group in the ventral lateral medulla and the A2 group in the nucleus of the solitary tract (Sawchenko and Swanson, 1982; Liposits et al., 1986; Cunningham and Sawchenko, 1988). When an animal is exposed to a threat, noradrenaline levels within the PVN are elevated (Pacak et al., 1992) and lesioning these noradrenergic pathways reduces corticosterone responses to stress (Ritter et al., 2003; Flak et al., 2014). In addition to innervating the PVN, noradrenergic fibers also innervate the median eminence (Palkovits et al., 1980). No past studies have investigated whether noradrenaline regulates CRH nerve terminal function in the median eminence. Contrary to our expectations, noradrenaline induced a robust inhibition of action potential dependent Ca^2+^ elevations and suppressed CRH secretion. This inhibitory effect was dependent on the activation of α2-adrenoreceptors because it was prevented by prior incubation of brain slices with yohimbine. Our previous work using GCaMP Ca^2+^ imaging in CRH neuron cell bodies demonstrated that noradrenaline induces an excitation that depends on α1-adrenoreceptors (Gouws et al., 2022). However, this work also showed that α2-adrenoreceptors are also present on the soma/dendrites of CRH neurons and limit the extent of α1-mediated excitation.

It is currently unknown whether the noradrenergic fibers that innervate the median eminence arise from the same population of noradrenergic neurons that innervate CRH cell bodies in the PVN. This gap in our knowledge leaves open several possibilities as to what role median eminence noradrenaline might be playing in stress axis control. If noradrenergic fibers originate from a single population of neurons, then it is possible that noradrenaline may act to limit CRH secretion into the blood during times of stress and high CRH neuronal spiking activity. This mechanism would still allow high levels of CRH release at other central targets that are not innervated by noradrenergic afferents. If, on the other hand, noradrenergic fibers in the median eminence originate from a population of noradrenaline neurons distinct from those which innervate the PVN, then this would allow these different neural populations unique control of CRH secretion in response to different physiological/behavioural states. Finally, it is also possible that noradrenaline circulating in the blood (released from the adrenal medulla) may act on CRH nerve terminals. The median eminence has a leaky blood brain barrier and median eminence nerve terminals are exposed to bloodborne molecules. This mechanism may be one way in which HPA axis activity could be modulated based on the level of sympathetic output from the adrenal medulla.

In this current study, we have also shown that corticosterone can inhibit CRH peptide secretion directly from median eminence nerve terminals. For these experiments, corticosterone was present on the brain slice for 20 minutes before testing for an effect. The inhibition was also prevented with prior incubation of brain slices with the glucocorticoid receptor antagonist, RU486. The time course of the inhibition suggests that corticosterone is inhibiting secretion via a non-genomic glucocorticoid receptor pathway. This makes sense given that CRH nerve terminals lack a nucleus and therefore it would not be possible for glucocorticoid receptors to signal via their classical (slow) genomic action. Interestingly, we did not find any evidence of corticosterone inhibiting nerve terminal Ca^2+^ elevations. This suggests that the inhibitory effect may be mediated via regulation of vesicle release machinery as has been suggested for other neuromodulators (Iremonger and Bains, 2009).

This current data establishes the median eminence as a key site of glucocorticoid negative feedback in the brain. Past work has reported glucocorticoid binding sites in the rat median eminence (Sarrieau et al., 1988). However, the effect of glucocorticoids on CRH secretion from nerve terminals in past studies has been mixed. One study using synaptosomes isolated from the sheep hypothalamus demonstrated a suppression of electrically evoked CRH secretion following 40-60 min of corticosteroid treatment (Edwardson and Bennett, 1974). However, a study in rat tissue found that the glucocorticoid receptor agonist dexamethasone had no effect on basal CRH secretion from the median eminence (Suda et al., 1985). Of note, the effect of dexamethasone on stimulation-evoked CRH release was not tested in this last study.

Fast corticosteroid negative feedback of the HPA axis has been known to exist for many years (Keller-Wood and Dallman, 1984; Dallman, 2005; Andrews et al., 2012; Deng et al., 2015; Osterlund et al., 2016). In vitro studies have suggested that fast negative feedback could be mediated in part via activation of glucocorticoid receptors on CRH neuron cell bodies in the PVN (Di et al., 2003; Nahar et al., 2015). However, in vivo fiber photometry recordings of PVN CRH neuron activity in freely behaving mice have recently revealed that corticosteroid negative feedback of excitability only starts to manifest 30-40 min following elevation of corticosteroid levels. This study however did reveal that corticosteroid inhibition of pituitary ACTH secretion was much faster (Kim et al., 2019b). While direct inhibition of pituitary ACTH release has been clearly demonstrated (Keller-Wood and Dallman, 1984; Deng et al., 2015), our current data suggest that corticosteroids can also act directly on median eminence nerve terminals to quickly suppress CRH secretion. This could contribute to the fast suppression of ACTH in response to elevated levels of corticosteroids.

It is likely that CRH nerve terminals are also subject to regulation by other neurotransmitters, neuromodulators and hormones. Indeed, past work has shown that GABA can directly excite CRH nerve terminals in the median eminence (Kakizawa et al., 2016). Nerve terminal regulation is not limited to CRH neurons, as the excitability of gonadotropin-releasing hormone (GnRH) nerve terminals in the median eminence is also directly regulated by neuropeptides (Glanowska and Moenter, 2015; Iremonger et al., 2017). And more broadly, nerve terminal regulation has been shown to be a common mechanism across a wide variety of neurons in the brain (Nadim and Bucher, 2014).

In conclusion, this current study demonstrates that CRH nerve terminals are an independent point of regulation for CRH secretion. We show for the first time that noradrenaline has a powerful inhibitory action on CRH nerve terminals. Uniquely, while noradrenaline is inhibitory at nerve terminals, it exerts the opposite actions at the CRH soma. Finally, we reveal that corticosterone also inhibits nerve terminal CRH secretion and identify median eminence nerve terminals as an important site of corticosteroid negative feedback in the brain. Together these findings suggest that CRH nerve terminals represent a critical point of regulation in the control of the stress axis.

## Acknowledgments

This work was funded by a Royal Society of New Zealand Marsden Fund grant. We would like to thank Dr. Kenichi Okamoto (University of Toronto) for providing the cAMP Förster resonance energy transfer (FRET) probe (CEY) cDNA construct, and Dr. Rithwik Ramachandran (Western University) for providing critical support for the development of the sniffer cells. Thanks also to Professor Colin Brown and Dr Michel Herde for feedback on an earlier version of this manuscript.

